# Flow-mediated self-organization of ventilation in honeybee nests

**DOI:** 10.1101/212100

**Authors:** Jacob M. Peters, Orit Peleg, L. Mahadevan

## Abstract

European honey bees (*Apis mellifera*) live in large congested nest cavities with a single opening that limits passive ventilation. These nests are actively ventilated by individual bees which fan their wings at the nest entrance when the local air temperature exceeds a threshold. Here we show that colonies with relatively large nest entrances use an emergent ventilation strategy where fanning bees self-organize to form fanning groups, separating regions of continuous inflow and outflow. The observed spatio-temporal patterns correlate the air velocity and air temperature along the entrances to the distribution of fanning bees. A mathematical model that couples these variables to known fanning behavior of individuals recapitulates their collective dynamics. Additionally, the model makes predictions about the temporal stability of the fanning group as a function of the temperature difference between the environment and the nest. Consistent with these predictions, we observe that the fanning groups drift, cling to the entrance boundaries, break-up and reform as the ambient temperature varies over a period of days. Overall, our study shows how honeybees use flow-mediated communication to self-organize into a steady-state in fluctuating environments.

---

Many animal groups are able to solve complex problems using the collective action of individuals with limited sensory and cognitive abilities. These problems often involve spatial scales that are orders of magnitude larger than individuals that typically can sense and respond only locally. Thus, collectively organized solutions to problems such as complex navigation [1], predator avoidance [2] and distributed foraging [3] arise from the active interactions between individuals, which allow locally-sourced information to be integrated by the group [4].

Social behaviors in insects provide many such examples such as foraging recruitment, nest site selection and nest building. The solutions are often organized by a process called stigmergy in which individuals interacting with a common environment can lead to an emergent scheme or pattern without direct interaction [5]. In classical examples of stigmergy, individuals deposit static cues in the environment such as pheromones, food stores or building materials that influence future behaviors by other individuals through positive feedback or recruitment which lead to the emergence of pheromone trails or architectural features such as walls, piles or pillars [4]. In less commonly studied class of behaviors, individuals interact with a dynamical physical process, such as flow, to structure their behavior. Experiments on corpse clustering by the ant *Messor sanctus* [6] and nest tube blocking in leaf-cutting ants *Acromyrmex ambigus* [7] have demonstrated that depositing materials in the environment can modify airflow which can in turn influence future depositions. Several theoretical studies have suggested that if an individual-level behavior is responsive to flow-mediated cues and such behaviors manipulate flow, global behaviors can arise that control this flow for adaptive functions such as ventilation or thermoregulation [8, 9, 10].

A particular instance of this local-global behavior is seen in the context of thermoregulation and ventilation in honeybee colonies (> 10, 000 bees) that live in congested enclosures such as tree hollows or other pre-existing cavities, where they face the continuous challenge of maintaining relatively stable temperatures (≈ 36°C) and respiratory gas concentrations [11, 12]. Active ventilation is a natural solution to both problems as it circumvents the limits imposed by impervious walls, small entrances (relative to nest volume) and large colony size. This allows for colony-level gas exchange with the environment and prevents buildup of heat and CO_2_ within the nest [13, 14]. Indeed, groups of honey bees orient themselves with their abdomens facing away from the nest and actively pull air out of the nest by fanning their wings at the nest entrance. However, conservation of mass requires that air drawn from the nest must be balanced by air flowing into the nest. Southwick and Moritz [15] observed that colonies in hives with small round entrances (2cm diameter, 3.14cm^2^ area) exhibit tidal ventilation in which honey-bees actively draw air out of the nest entrance for a while and then stop, allowing air to passively flow back into the nest. They also suggested that bees might select nest sites with larger entrances to allow for unidirectional or bidirectional air flow. Since nest entrances in feral colonies have a range of shapes and sizes that spans more than an order of magnitude [11], a natural question is if and how the mechanism described in [15], i.e. temporal modulation of in/outflow, generalizes to characterize ventilation dynamics in nests with moderate to large entrances.

To answer this question, we quantify the fanning behavior of bees at a large nest entrance shown schematically in Fig. 1A. We used four Langstroth beehives (80 liters, 20,000-40,000 bees each) with a single slit-like rectangular nest entrance (2×36cm) that we monitored over time. In order to quantify the influence of the distribution of fanning bees along the entrance on the induced flow pattern, we counted the number of fanning bees in each of fifteen bins along the entrance (see Fig. 1D and SI Movie S3). Although there are fanning bees just inside the entrance, we counted only the visible ones, which serve as a proxy for local fanning intensity. Simultaneously, we used a directional VelociCalc anemometer to measure the flow speed perpendicular to the entrance at the boundaries between the bins, with flow being positive outwards, and negative inwards. Flow direction was measured by placing a wool fiber at each position; if its motion was imperceptible or non-directional, no velocity value was recorded (see SI Movie S1). A thermocouple attached to the tip of the anemometer allowed us to measure the temperature at each location, which can also be qualitatively visualized using an infrared camera (Figure 1B). All measurements were carried out five times a day over the course of three consecutive days. In Figure 1D, we show the density of fanning bees, air velocity and air temperature as a function of the position along the entrance for one of the hives observed (see SI Fig. S1 and Fig. S2 - for the complete data set involving multiple bee hives).

**Figure 1:**
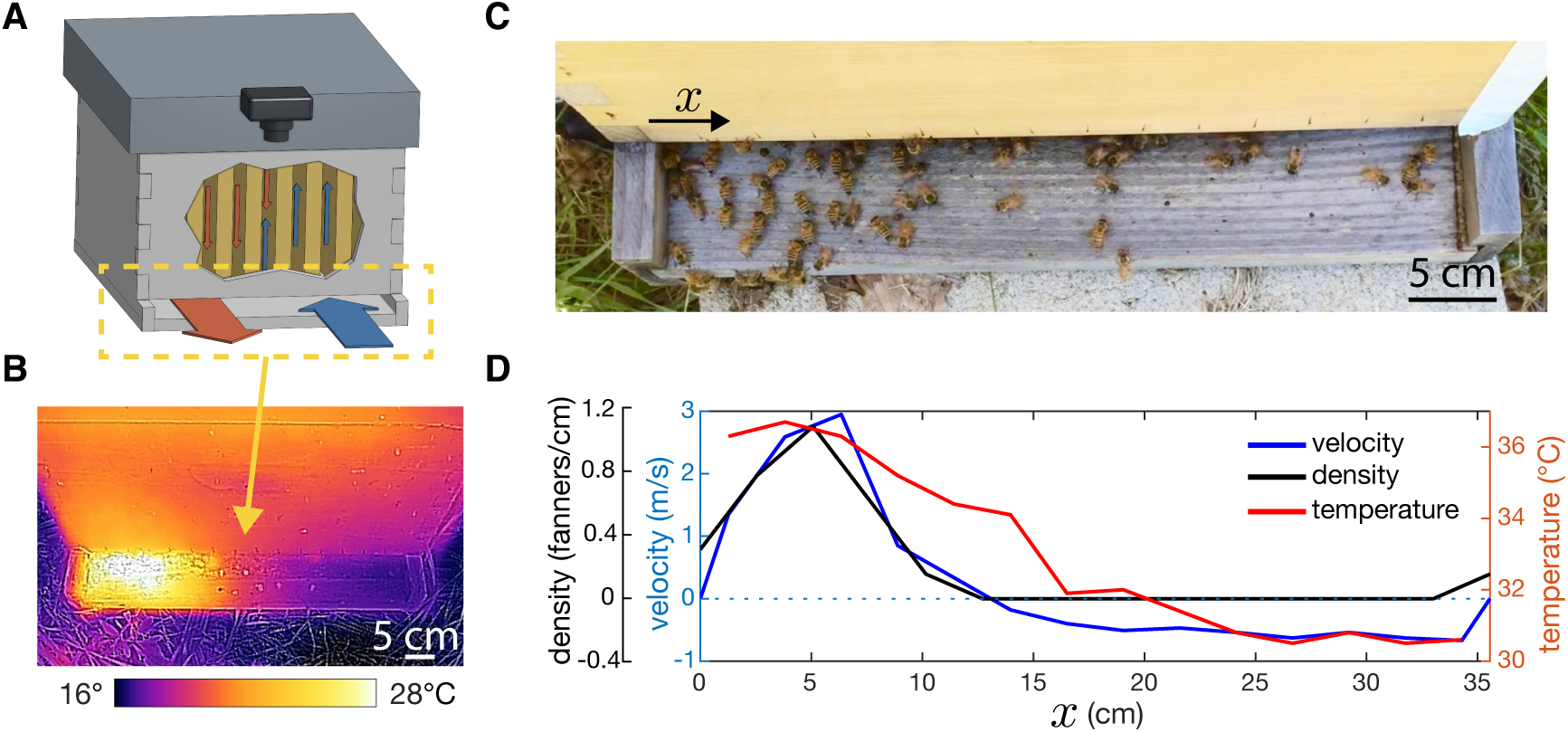
A) A schematic illustrating the path of air through the hive as induced by fanning bees. Warm, stale air is drawn out of the entrance by fanning bees and relatively cool ambient air enters passively where fanning bees are absent. The camera indicates the viewing angle in B and C. B) A thermal image of the hive entrance at night when fanning bees are actively fanning. Notice inflowing air at the right of the entrance has cooled the wood and the outflowing air induced by fanning bees on the left has warmed the wood. C) Honeybees ventilating at the entrance of a hive. Note the dense group of fanning bees at the left of the entrance and the lack of fanning bees at the right of the entrance. D) The air velocity (blue) and temperature (red) along the nest entrance of a hive. Note that inflow is indicated by negative values and outflow is indicated by positive values. These data demonstrate that the temperature profile along the entrance can be used as a qualitative proxy for flow velocity.

We observed correlated variations in density, velocity and temperature at the entrance across space and time. In contrast with the observed rapid temporal modulation of ventilation behavior in nests with small openings [15], our observations show that for larger entrances, ventilation behavior is spatially modulated (i.e., in/outflow separated in space) but temporally steady, at least over times when the ambient temperature was steady. This dynamic adaptation to the physiological needs of a colony demands a dynamic explanation that links the behavior of individuals distributed at the hive entrance to the observed correlations between fanner density, air velocity, and air temperature in space and time.

Honeybee colonies show broad inter-individual variation in the temperature thresholds that induce fanning, in part because of their high genetic diversity [16, 17]. This variation can lead to emergent task allocation via the so-called task threshold model [18], which states that when the demand for a task is large, more individual bees will respond due to the broad variation in the task thresholds [17]. This variation is higher in colonies with a queen which has mated multiple times [16], and promotes the temporal stability of thermoregulation. Furthermore, honeybees that are heated in a laboratory setting are more likely to fan at a given temperature when they are in a group than when they are alone [17]; independent of group size, individuals showed broad variation in temperature thresholds. In the largest group size considered (10 individuals), the mean temperature at which bees begin to fan was near the preferred hive (i.e., brood) temperature (≈ 36°C).

Our minimal framework for the spatiotemporal organization of fanning starts by characterizing the local fanning response of individual bees to the local air temperature. We must also account for fluid flow which is induced by the bees and which carries the signal to which the bees are responding (i.e., heat derived from the hive). For simplicity, we consider the case when the hive and environmental temperatures are constant and focus on the dynamics of the bees at the entrance.

To link bee behavior, air temperature and airflow we need to quantify how (i) the distribution of fanning bees, *ρ*(*x, t*), (ii) the local air temperature, *T* (*x, t*), and (iii) the local flow velocity, *v*(*x, t*) vary with time *t* along the nest entrance. Because the nest entrance in this hive has a high aspect ratio (short but wide) we can model the entrance *x* as a 1D line (see Figure 2A).

**Figure 2:**
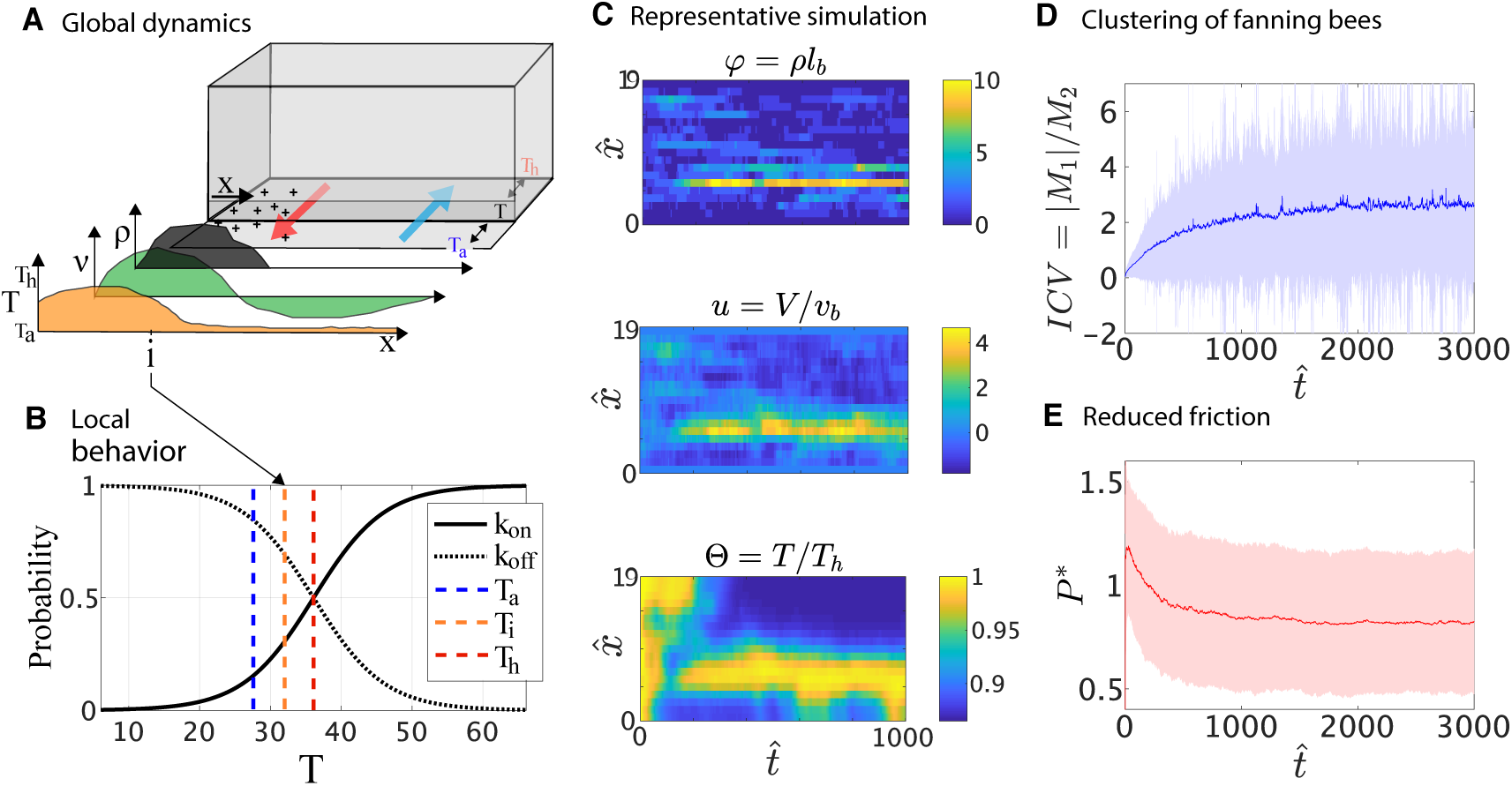
Theoretical model and numerical simulations. (A) Nest entrance represented as a one-dimensional line (*x*). Our model relates the density of fanners (*ρ*(*x, t*)), air velocity (*v*(*x, t*)), and air temperature (*T* (*x, t*)). Air drawn from the entrance by the bees has a +ve velocity. (B) The probability that a bee will begin fanning at position *x* and time *t* (*k*_on_) is high at high temperatures and low at low temperatures. Conversely, the probability that a fanning bee will cease fanning (*k*_off_) is high at low temperatures and low at high temperatures. (C) The density of fanning bees *ρ*(*x, t*), (D) local air temperature *V* (*x, t*) and (E) the local air velocity over the first 1000 steps of a representative numerical simulation. The distribution of fanners was initially uniform, the temperature was initially 36°C along the entire entrance and the velocity was initially 0. Ambient temperature was fixed at 28°C. (F) The mean inverted coefficient of variation (*ICV*) is plotted for the first 3,000 time steps of 1000 simulations. Error bands indicate standard deviation. High *ICV* indicates that fanners are highly clustered. (G) The power *P* ^∗^ lost to friction throughout the simulation. Note that *P* ^∗^ is a scaled or dimensionless metric and has no units. As fanners become more clustered, the amount of fluid friction is reduced, indicating that self-organization leads to increased ventilation efficiency.

Consistent with the task threshold model, we assume that the probability of a bee stopping and initiating fanning behavior is determined by the local temperature [17]. Therefore, the local density of fanning bees, *ρ*(*x, t*) changes according to the equation

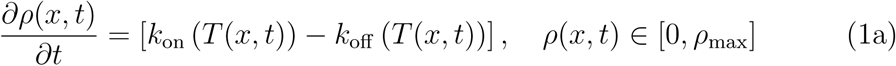

where *k*_on_ is the rate at which bees initiate fanning behavior, *k*_off_ is the rate at which they cease fanning behavior, and *ρ*_max_ is the maximum density achievable (given spatial constraints at the nest entrance). These rates are assumed to be sigmoidal functions of the local air temperature, i.e.

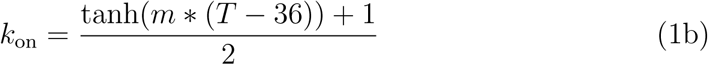

and

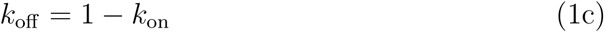

where *m* controls the slope of the sinusoidal function (see Figure 2B), and is fit to approximately reproduce the variation in observed temperature thresholds [17] (see S4.1). Although, recent studies of fanning behavior in controlled laboratory settings suggest that the thermal response thresholds for fanning are affected by group size [17], presence/absence of larvae [19], and heating rate [20], our minimal representation of the fanning response as a switch-like behavior allows us to focus on the interaction with airflow and temperature.

To characterize the air flow, we assume that each bee generates an outward air flow with velocity *v*_*b*_. Because the nest has just one opening, air that is actively drawn from the the entrance must be balanced by air flowing passively into the entrance elsewhere in order to ensure conservation of mass. Flow conservation at the entrance demands that (see SI for a simple derivation of this relation)

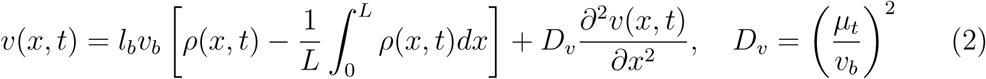

where *L* is the size of the nest entrance, *l*_*b*_ is the wingspan of a bee, and *µ*_*t*_ is the turbulent viscosity of air. The first term characterizes the difference in the local density of fanning bees from the average density over the entire length of the entrance. This term conserves the volume of air in the hive (since the net flow rate vanishes). The second term is associated with fluid friction with *D*_*v*_ being an effective kinematic viscosity.

Finally, to characterize the dynamics of the local air temperature along the entrance, we assume that air temperature is governed by the local velocity and temperature difference between the entrance temperature and the upstream temperature, and can be described by a modification of Newton’s law of cooling (neglecting complex flow dependences, see SI):

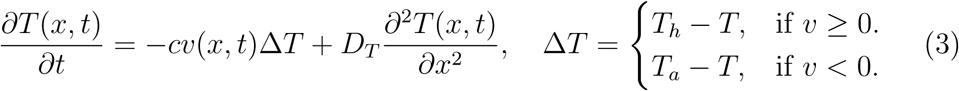

where *c* is a characteristic inverse length (chosen so that Newtonian cooling due to fanning dominates lateral diffusion), where *T*_*a*_, *T*_*h*_ are the ambient and hive temperature, and *D*_*T*_ is the thermal diffusivity.

The variables in our model can be rescaled using the following definitions: 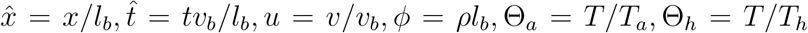, which leads to a dimensionless set of our original equations with four dimensionless parameters: (i) *L/l_b_*, a dimensionless measure of the entrance length (ii) 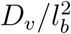, a measure of the effect of fluid friction, (iii) *D_T_ /v_b_*, a scaled heat diffusion coefficient and (iv) *l_b_/v_b_*, a characteristic time scale associated with bee fanning. In addition, we have two parameters that characterize each of the sigmoids associated with the switching of the fanning response.

To complete the formulation of the model, we need to specify boundary conditions for the temperature and velocity of the airflow. The air velocity *v* is assumed to be zero at the ends of the nest entrance, while the temperature was assumed to satisfy the Robin boundary condition

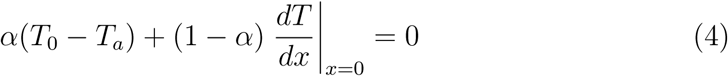

where *T*_0_ is the temperature at the boundary and *α* prescribes the thermal conductivity of the boundary (i.e., the walls on either side of the entrance). Then *α* = 0 corresponds to the boundary being a perfect insulator while *α* = 1 corresponds to the boundary being a perfect conductor (see Fig. S5).

We used MATLAB to solve the initial boundary value problem (1-4) using a finite difference scheme in space and a Runge-Kutta method in time, using the following values for the parameters: width of the nest entrance *L* = 0.38m, wingspan of a bee *l*_*b*_ = 0.02m, air velocity generated by an individual fanning bee *v*_*b*_ = 1m/s [21]. The diffusion coefficients *D*_*v*_ and *D*_*T*_ were fit to match observed behavior (*D*_*v*_ = 1 × 10^*−*4^ and *D*_*T*_ = 4 × 10^−5^; see Figure S3 for a sweep of these parameters). All simulations were executed with the following initial conditions: 1) Fanning bee density initially given a uniform distribution with one fanning bee per bin (1 bin = *l*_*b*_), 2) local air velocity was zero along the length of the entrance, and the local air temperature was initially *T*_*h*_.

Fanning bees initially formed multiple clusters allowing for spatial separation of inflow and outflow (Figure 2c). Over time, several dominant clusters grew as other smaller clusters petered out. By 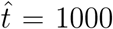 (≈ several minutes), a single dominant cluster emerged leading to one region of outflow and 1-2 regions of inflow. This condition appeared to be stable, however the position of the fanning group varied across iterations. We quantified this clustering of fanning bees using the inverse of the coefficient of variation of the density of fanning bees, i.e. *ICV* = |*M*_1_| */M*_2_, where *M*_*i*_ is the *i*^*th*^ moment of the density. If ICV is large, the bees are highly clustered. We also quantified the amount of power lost to fluid friction using the dimensionless parameter 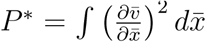. As shown in Figures 2F and 2G, clustering of fanners was inversely related to the amount of fluid friction in the system, suggesting that self-organization leads to more efficient ventilation by reducing friction (or shear) at the nest entrance.

When the simulations were run over long times, 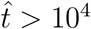 (*≈* several hours), the dominant cluster drifted in space (Figure 3A). When *T*_*a*_ was much less than *T*_*h*_ (∆*T* 10°C), the cluster drifted relatively freely. However, at higher *T*_*a*_ the cluster tended to cling to the boundaries of the entrance with occasional spontaneous switching from side to side. As *T*_*a*_ approached *T*_*h*_ (∆*T* 2°C), the dominant cluster broke up to form multiple ephemeral clusters. Our model demonstrates that the interaction between fanning bees and the flow field in which they are embedded allows for emergent clustering of fanning bees and ultimately leads to ordered, efficient airflow through the nest entrance without a leader or central coordination. An explicit prediction of the model is that the dominant cluster should drift in space over long time scales and that this drift is qualitatively different at various temperatures.

**Figure 3:**
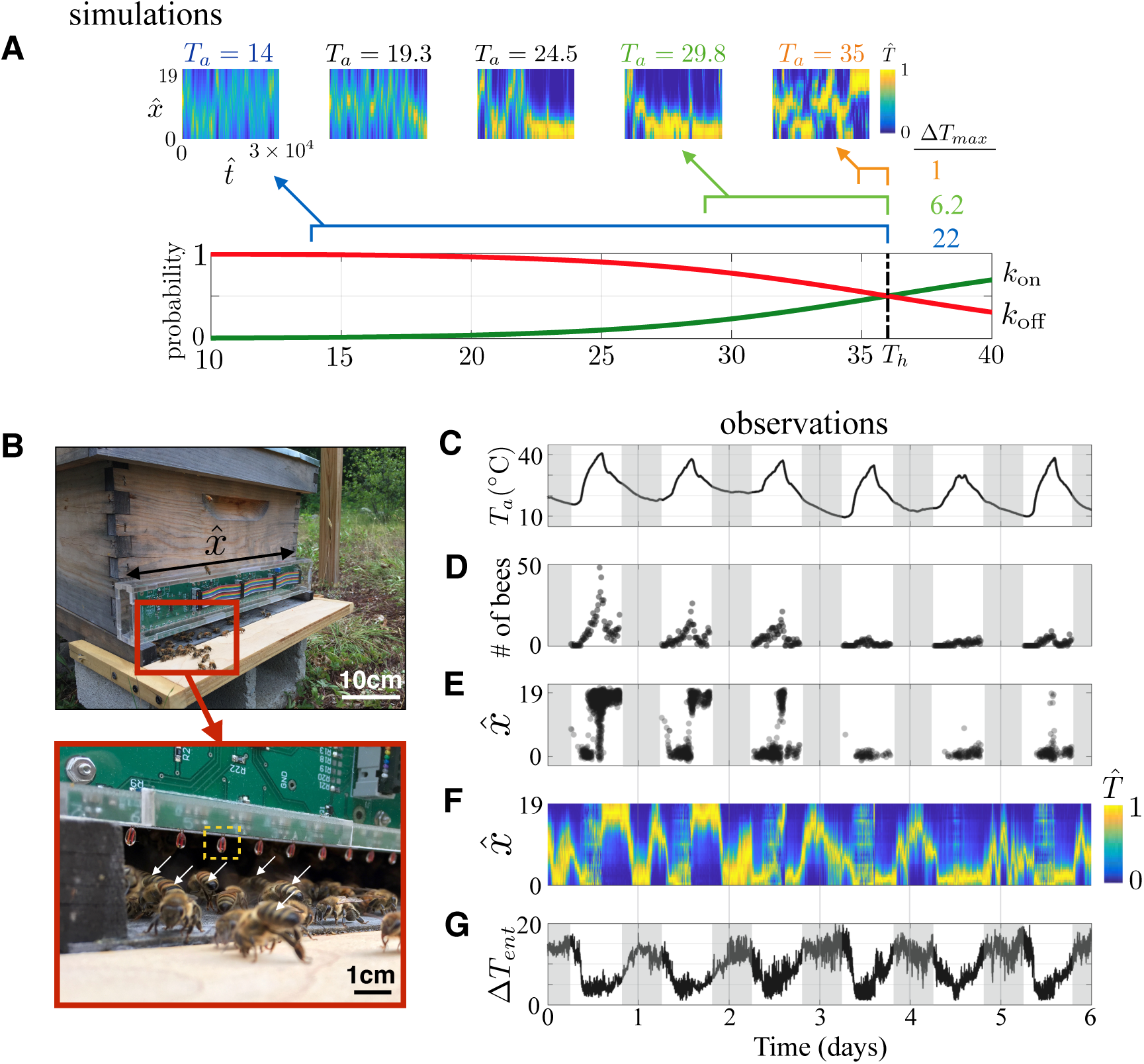
Predicted and observed ventilation dynamics over long time scales. A) Numerical simulations predict that the position of the fanning group will drift over long time scales (30,000 time steps, *T*_*h*_ = 36°C, *D*_*v*_ = 1*e^−^*^4^, *D*_*t*_ = 4*e^−^*^5^, *m* = 0.1, *c* = 0.05, *α* = 0.2). At low temperatures (*T*_*a*_ = 15, 20.3°C), the fanners tend to occupy the center of the nest entrance and drift in space. At higher temperatures (*T*_*a*_ = 25.5, 30.8°C) the fanning group clings to the boundary of the nest entrance. When ambient temperature approaches hive temperature (*T*_*a*_ = 36°C), no singular, persistent fanning group emerges. (B) The temperature sensors along the entrance allow a true measure of air temperature in the flow stream. Fanning bees are indicated with white arrows. (C) The diurnal oscillations in ambient temperature closely tracked fanning intensity. (D) Total fanning bee number over time - gray regions indicate dark hours when it was difficult to record fanning behavior directly. (E) The position of the fanning bees indicates that single fanning group forms except when *T*_*a*_ is very close to *T*_*h*_. During warm hours the fanning group tends to cling to the boundaries of the nest entrance. (F) The local air temperature shows that the position of the fanners is associated with warm, out-flowing air. The position of the fanning group (and outflow) tends to drift during the night when *T*_*a*_ is low. The entrance temperature has been normalized 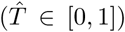. (G) The range of local air temperatures measured at any instance in time (∆*T*_*ent*_ = *T_max_ − T_min_*) is also reported.

In order to test these predictions, we developed a method to continuously monitor the position of fanning bees at the nest entrance and the resultant flow pattern in the field under naturally varying conditions over many days. Although no appropriate technology exists to continuously measure air velocity simultaneously at many positions within a cluttered and dynamic environment, we were able to use temperature as a qualitative proxy for flow direction at the nest entrance (see Fig. 1D). We placed custom 32-sensor arrays above the nest entrances (1.8 × 37cm) of three additional Langstroth beehives enabling us to sample local air temperature with high spatial and temporal resolution (1.17cm sensor spacing, 10 second sampling intervals, Figure 3B). Short videos were taken of the entrance of one of these hives every 10 minutes during daylight hours and the position of visible fanning bees was recorded (SI Movie 2-3). See Section S3 for details.

The ambient temperature oscillated according to a 24 hour diurnal cycle (Figure 3C) with variation in minimum and maximum daily temperatures throughout the observation period. The total number of fanning bees visible at the nest entrance tracked these oscillations and fanning intensity was higher on warmer days (Figure 3D). The position of fanning bees during warm daylight hours revealed that the fanners tend to form well-defined clusters which tend to cling to the boundaries of the nest entrance (Figure 3E). On warmer days, the ambient temperature would approach the nest temperature at midday and the fanning group would brake up into multiple, less-defined clusters. When the ambient temperature fell again, a single cluster would again emerge. For the hive depicted in Figure 3 (Hive 1), the dominant fanning group often occupied the east side of the entrance during the morning and the west side of the entrance during the afternoon. This suggests that under some conditions, solar radiation may impose an environmental asymmetry that can bias the position of the fanners. This was not the case for Hives 2 and 3, suggesting that this is not the only factor determining the position of the fanners. The air temperature profile along the nest entrance reflected the distribution of fanning bees. Figure 3F maps the normalized air temperature 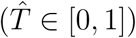 at each time step so that the entire dataset can be represented with one heat map. The difference between the minimum and maximum temperature at the nest entrance (∆*T*_*ent*_ = *T_max_ T_min_*) is also plotted in (Figure 3G). The position of the dominant fanning group as indicated by the warm, outflowing air tended to drift away from the boundaries during the night when ambient temperatures were low as predicted by the model.

The nuances of the ventilation behavior are inevitably affected by the particular environmental conditions experienced by a colony, and yet the qualitative predictions made by our model are born out in the observed behavior in naturally fluctuating conditions. When *T*_*a*_ is lower than *T*_*h*_ a single cluster of fanners forms and tends to drift in space. As *T*_*a*_ increases the cluster tends to fix to a boundary. Which boundary the cluster fixes to may be biased by asymmetries in the environment. Finally, as *T*_*a*_ approaches *T*_*h*_, the cluster breaks up into multiple or less defined clusters. Our observations are in agreement with theoretical predictions and suggest that collective nest ventilation is not just a product of the bee behavior, but arise from the local flow-mediated interactions between individual bees and of the resultant hive-scale fluid dynamics.

There are two behavioral components of this process that are critical to self-organized ventilation. First, the bees must (and do) fan air out of rather than into the nest entrance. This allows the bees to sense the upstream nest temperature. If the bees fanned into the nest entrance, they would have no information about the state of the hive. Interestingly, another cavity nesting honeybee species, *Apis cerana*, fans into the nest entrance [22]. This species likely uses an alternate strategy to the one described here or occupies nests with a small nest entrance in which spatial organization is not required [15]. Second, the switch function which determines the probability at which a bee will fan at a given temperature, and has likely been tuned through natural selection. If the slope of this function is too shallow, fanning behavior is weakly coupled to temperature and no organization will emerge (see Fig. S5A,E). If the slope is too steep, fanning behavior can occur only over a small range of temperatures (see Fig. S5C,G). Indeed, colonies with high genetic diversity have more variation in individual temperature thresholds for fanning are able to achieve a more stable hive temperature through time [16]. We have shown that this diversity is also critical to the stability of spatial patterning of fanning behavior which is required for efficient ventilation.

Our study demonstrates how harnessing the dynamics of the physical environment allows for large scale organization of a physiological process. This differs from classical stigmergy, which facilitates coordination by integrating spatially static information over longer time scales. Honeybees sense local air temperature (which is coupled to speed and direction of airflow) and drive airflow when temperatures are high. Because the individuals are embedded in a common flow-field, their behavior is influenced by nonlocal interactions mediated by flow. The selforganization of fanners into groups which efficiently partition inflow and outflow, reduce friction and avoid antagonistic fanning behavior is ultimately the result of flow-mediated information processing that integrates locally sourced information over large spatial scales even in the absence of direct interaction between neighboring individuals. This ability to manipulate existing physical processes locally to create self-organized behavior on large scales may be a pervasive strategy in the evolution of complex systems.

## Acknowledgments

We thank Jim MacArthur for building the sensor array and other valuable advice on instrumentation, Andrew Clark for assistance with hive monitoring, Tom Seeley and Michael Smith for discussions and guidance, and an NSF GRFP DGE1144152 (JMP), and NSF PHY1606895 (OP,LM) for partial financial support.

